# Sexually selected traits predict female-biased philopatry in a sex-role reversed shorebird

**DOI:** 10.1101/2025.01.13.632832

**Authors:** Leilton W. Luna, Sara E. Lipshutz

## Abstract

Sex-biased dispersal plays a key role in shaping population dynamics and genetic structure. Two main hypotheses have been proposed for how territoriality and mating competition impact sex-biased dispersal. Female-biased dispersal is expected in monogamous systems with male resource defense, whereas male-biased dispersal is expected in polygynous systems with male competition over mates. However, patterns of sex-biased dispersal in polyandrous species, where females compete for both territories and mates, remain poorly understood. We investigated sex-biased dispersal in two polyandrous Jacana species, the Northern Jacana (*Jacana spinosa*) and Wattled Jacana (*J. jacana*), which exhibit intense female-female competition for territories and mates and differ in the strength of sexual selection. We analyzed sex-biased dispersal by assessing genetic differentiation and individual assignment indices in these jacana species across Central America. Our findings reveal strong male-biased dispersal in Northern Jacanas, indicated by higher genetic structuring and philopatry in females. In contrast, Wattled Jacanas showed no significant dispersal bias between sexes. Furthermore, sexually selected traits in Northern Jacana females, such as larger body mass, tarsus, and wing spur length, were associated with philopatry, suggesting that larger females retain territories, whereas smaller females disperse. To our knowledge, this is the first genetic evidence of male-biased dispersal in a polyandrous species. Our findings reveal that sexually selected traits, in addition to territorial and mate competition, are important for understanding species and sex differences in dispersal evolution.

## Introduction

Sex-biased dispersal, where males and females exhibit different movement patterns, plays an essential role in population dynamics and genetic structure (Mishra et al., 2020; Cano et al., 2008). Traditionally, the primary drivers of sex-biased dispersal have been attributed to ecological and social factors, such as competition for resources and mates (Dobson, 1982; Greenwood, 1989; Perrin & Mazalov, 2000). These drivers are thought to differ between species based on mating systems and reproductive strategies (Trochet et al., 2016; Li & Kokko, 2019). The local resource competition hypothesis posits that female-biased dispersal is expected in socially monogamous systems where males defend resources (Greenwood, 1989). In contrast, the mate competition hypothesis predicts male-biased dispersal in polygynous systems where males compete for mates (Dobson, 1982; Devillard et al., 2004; Pérez-Espona et al., 2010). However, taxa with similar mating systems can show varying intensity and direction of sex bias in dispersal behavior, pointing to additional levels of complexity (Trochet et al., 2016; Pérez-González & Carranza, 2009).

An important, but often-overlooked factor in sex-biased dispersal is the investment in sexually selected traits, such as body size, ornaments, and weaponry. Dispersal tends to be limited in the sex bearing the higher maintenance costs of these traits, typically males (Lane & Shine, 2011; Trochet et al., 2016), while the sex with less costly traits may be more inclined to disperse. Costly secondary sexual traits impair dispersal by requiring substantial resources, increasing predation risk, and favoring philopatry, as staying local often provides better mating opportunities than dispersing (Badyaev & Duckworth, 2003; Mobley et al., 2018; Henshaw et al., 2022). Considering the role of secondary sexual trait investment, in addition to mating systems, may provide a more holistic understanding of how sex-biased dispersal can evolve.

Hypotheses for sex-biased dispersal have largely been applied to male-male competition in socially monogamous and polygynous mating systems (Hammond et al., 2005; Cano et al., 2008; Dubey et al., 2008; Hardouin et al., 2021), leaving the role of female-female competition relatively unexplored. Socially polyandrous (*i.e.*, sex-role reversed) mating systems, in which females compete for territories encompassing multiple males, provide a unique opportunity to disentangle drivers of sex-biased dispersal from potentially confounding effects of sex.

Polyandrous systems also feature females with more extreme sexually selected traits than males, including larger body size and weaponry. However, making a direct prediction for sex-biased dispersal in polyandrous systems is not straightforward, as females compete both for territories and mates, which complicates disentangling the expectations of competing hypotheses. In alignment with the mate competition hypothesis, intense female-female competition for mates may drive female-biased dispersal. Alternatively, greater female defense of territorial resources may drive male-biased dispersal. As a novel third hypothesis, we propose that higher female investment in sexually selected traits such as body size and weaponry may contribute to sex-biased dispersal.

Tropical shorebirds in the Jacanidae family, such as the Northern Jacana (*Jacana spinosa*) and Wattled Jacana (*J. jacana*), exhibit social polyandry and intense female-female competition (Jenni & Collier 1972; Stephens 1984; Lipshutz, 2017). Despite sharing a mating system, these sister species differ notably in the intensity of sexual selection. Compared to Wattled Jacanas, Northern Jacanas show greater levels of social polyandry, with female Northern Jacanas defending more males than female Wattled Jacanas (Jenni & Collier 1972, Emlen et al 1998). Northern Jacanas also have more pronounced female-biased sexual dimorphism in sexually selected traits including body mass, tarsus length, and wing spur length, as well as higher levels of aggression (Lipshutz, 2017), and longer sperm (Lipshutz et al., 2023). These interspecific differences highlight how varying levels of sexual pressures on females can shape key behavioral and morphological traits. We hypothesize that the stronger sexual selection observed in Northern Jacanas drives pronounced sex-biased dispersal, while the comparatively weaker sexual selection in Wattled Jacanas results in more moderate or less evident dispersal patterns.

To test these hypotheses, we analyzed sex-biased dispersal in Northern and Wattled Jacanas by comparing genetic differentiation and assignment between males and females across localities in Costa Rica and Panama (Figure 1a). Genetic methods can indirectly infer sex-specific dispersal, as gene flow results from dispersal followed by breeding (Goudet et al., 2002). When the dispersal rate of one sex is relatively high and/or exhibits strong spatial autocorrelation, dispersal bias can be detected using biparentally inherited genetic markers (Goudet et al., 2002; Banks & Peakall, 2012). We examined genetic patterns at both population and individual levels. At the population level, we used the fixation index (*F*_ST_) to measure genetic differentiation among populations. At the individual level, we applied the assignment index (AIc) to estimate the likelihood that an individual’s genotype originated from a specific locality, distinguishing philopatric individuals from recent immigrants. We predicted that the dispersing sex would exhibit lower geographic genetic structuring due to higher gene flow, with lower *F*_ST_ indicating less genetic differentiation and negative mean AIc values, indicating higher frequency of recent immigrants. Finally, to assess whether larger sexually selected traits predict movement patterns, we compared morphological traits associated with reproductive success, including body mass, tarsus, and wing spur length (a weapon), between likely recent immigrants and philopatric individuals. We anticipated that recent immigrants would have less developed secondary sexual traits, suggesting that less competitive individuals are more likely to disperse.

**Figure 1.**
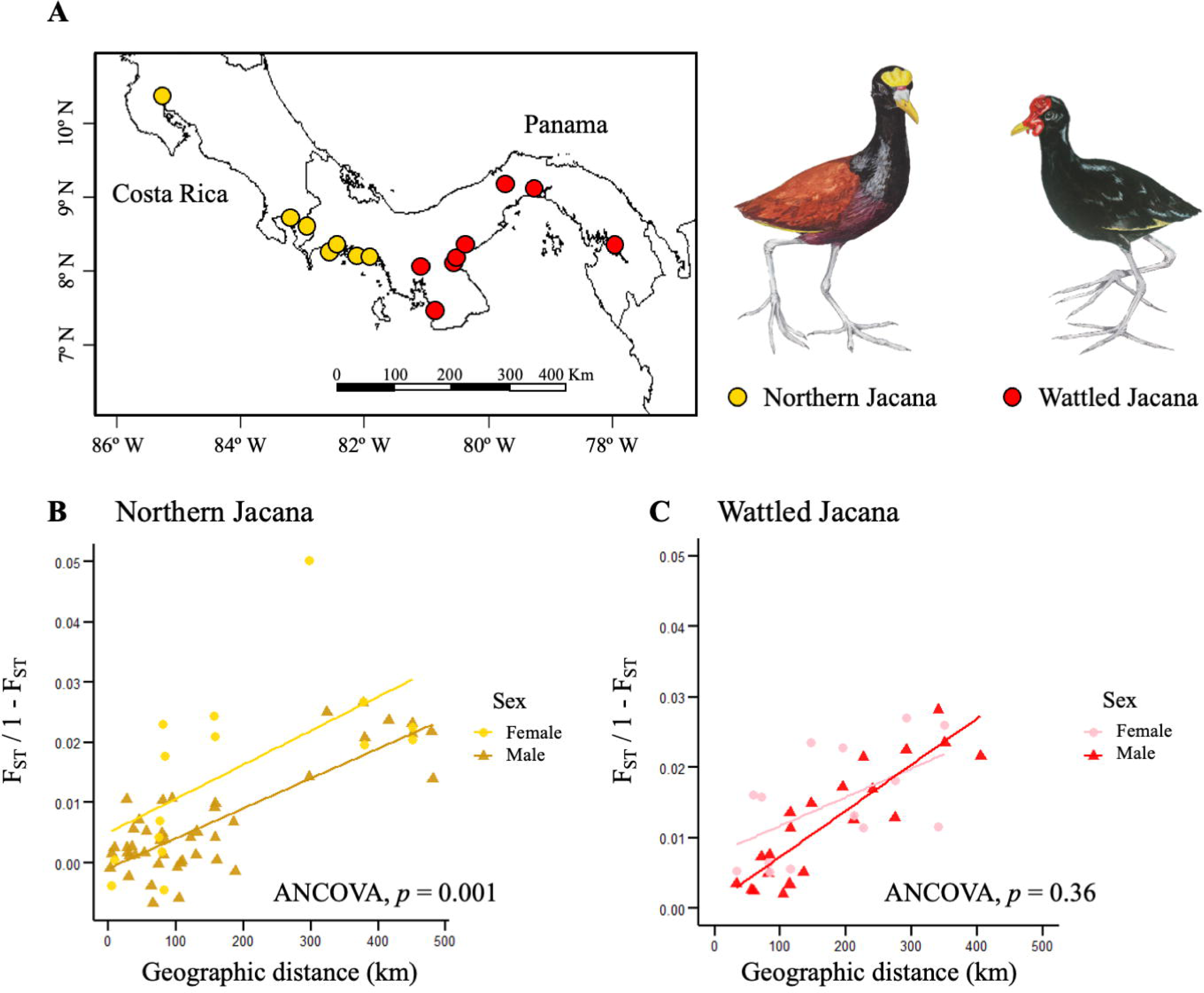
Sex-biased dispersal inferred from genetic differentiation between males and females of Northern Jacanas and Wattled Jacanas. (A) Geographic distribution of sampled localities in populations from Costa Rica and Panama. (B) Genome-wide differentiation (Weir and Cockerham’s weighted *F*_ST_) between sampled localities, measured in kilometers, comparing males and females of Northern Jacanas and (C) Wattled Jacanas.

## Materials and methods

### Sampling and genomics data processing

We used genotype-by-sequencing for 271 samples of Northern Jacana and Wattled Jacana, as published in Lipshutz et al. 2019 (BioProject PRJNA494934 and BioSamples SAMN10185940-SAMN10186205). Sampling included 15 localities across Costa Rica and Panama, with 7 localities for Northern Jacana and 8 for wattled jacana (Figure 1, Supplementary Information Table S1). The raw sequences were filtered to discard barcodes, adapter tags, and reads with a quality score below 20 using the process_radtags module in Stacks v2.68 (Rochette et al., 2019). Cleaned reads were indexed and aligned to a newly developed Northern Jacana Hi-C reference genome (Paton et al., in prep.) using BWA v0.7.18 (Li & Durbin, 2009) and processed with Samtools v1.21 (Li et al., 2009). We used the gstacks module in Stacks v2.68 to generate a catalog of consensus loci and SNPs. From the SNP catalog, we created unfiltered VCF files for the entire dataset, combining both species and potential hybrids using the *populations* module in Stacks v2.68 (Rochette et al., 2019). After identifying and excluding hybrid individuals (see below), we generated multiple VCF files for each Jacana species and sex, excluding scaffolds linked to sex chromosomes. For downstream analyses, we filtered our SNP dataset in VCFtools v.4.2 (Danecek et al., 2011) to include a minor allele frequency > 0.05, linkage disequilibrium < 0.1, removal of indels, and a maximum of 10% missing data. Due the reduced genomic distribution of this dataset, we identified limited uniparental inheritance markers, i.e. SNPs linked to the W-chromosome and mitochondria, that would facilitate evaluations of sex-biased dispersal. We found that no scaffolds were mapped to mitochondrial DNA, and as few as 6 SNPs were identified across the short scaffolds linked to the W-chromosome.

### Hybrid identification and exclusion

Northern and Wattled jacanas hybridize where their ranges overlap in western Panama (Lipshutz et al. 2019). Hybrids, with their mixed genetic profiles, may not accurately reflect the genetic structure of the parental populations, potentially biasing the analysis and leading to incorrect interpretations of isolation by distance between localities. To avoid bias introduced by hybrid individuals when estimating genetic differentiation metrics, we excluded first-generation (F1) hybrids and backcrosses. Hybrids were identified using two approaches: estimating admixture proportion (Q) and hybrid indices, with the programs STRUCTURE v2.3.4 (Pritchard et al., 2000) and the R package triangulaR v0.0.0.9 (Wiens & Colella, 2024), respectively. In STRUCTURE, we performed 10 replicates with K = 2, using a burn-in of 100,000 followed by 500,000 Monte Carlo iterations, and a mixture model with correlated allele frequencies. We used CLUMPP v1.1.2 (Jakobsson and Rosenberg 2007) to account for possible multimodality and label swapping between replicates, and Distruct2.py (Rosenberg 2004; Raj et al. 2014) to visualize Q values, classifying individuals as putative hybrids if 0.1 < Q < 0.9 (Supplemental Material Figure S1). In triangulaR v0.0.0.9, we estimated the hybrid index and interclass heterozygosity under Hardy-Weinberg equilibrium to classify F1 and backcrosses. Putative hybrid individuals with a hybrid index > 0.2 or < 0.8 and interclass heterozygosity > 0.25 were excluded from subsequent analyses. This process refined our sample set to 115 Northern Jacanas and 103 Wattled Jacanas (see Supplemental Material Figure S2 and Supplemental Information Table S1).

### Sex identification

Sex was classified in the field based on body mass distributions, following the methods outlined by Wrege and Emlen (2005). Ambiguous cases were further resolved using polymerase chain reaction (PCR) by sequencing the CHD region of the Z and W chromosomes, as detailed in Lipshutz et al. (2017). Additionally, we employed a joint sex determination framework based on the depth of coverage of sex-linked scaffolds using the R package SATC (Nursyifa et al., 2022). The depth of coverage of each sample read was mapped to scaffolds from the Northern Jacana reference genome, and depth of coverage values were normalized to distinguish heterogametic ZW females and homogametic ZZ males from the sex-linked scaffolds (Supplementary Information Table S2).

### Isolation by distance and genetic differentiation

To investigate sex and species-specific patterns of genetic differentiation across sampled localities, we estimated isolation by distance from correlation analyses of genetic differences and geographic distances separately for each group. Genome-wide differentiation for each dataset was estimated using Weir and Cockerham’s weighted *F*_ST_ between pairs of sampled localities, with a minimum of three samples per locality, using the R package hierfstat v0.5-11 (Goudet, 2005). Weir and Cockerham’s weighted *F*_ST_ corrects for biases in allele frequency estimation, especially in small sample sizes, and is effective for comparisons at different spatial scales (Weir & Cockerham, 1984). Pairwise Euclidean distances between localities were computed using the ’distm’ function in the R package geosphere v1.5-20 (Hijmans, 2024). Isolation by distance patterns were assessed with a Mantel test, employing 9,999 replicates, in the R package adegenet v.1.3-1 (Jombart & Ahmed, 2011). To evaluate whether the slopes of Mantel regressions differed between sexes within each species, we applied ANCOVA tests in R. Additionally, to assess overall genetic differentiation, we compared mean *F*_ST_ values between the two species and between the sexes within each species using a non-parametric T-test.

### Assignment indices

To assess sex-biased dispersal, we used the corrected assignment index (AIc), which indicates the likelihood of assigning an adult individual to a particular locality (Favre et al., 1997; Goudet et al., 2002). Negative AIc values signify potential recent migrants, while positive values denote residents (Favre et al., 1997; Goudet et al., 2002; Colson et al., 2013). Since immigrant individuals typically have lower AIc values than residents (Goudet et al., 2002), we predicted that the sex exhibiting greater dispersal would, on average, display lower AIc values than the more philopatric sex. AIc values were computed using hierfstat v0.5-11 (Goudet, 2005), and a permutation test was performed to assess differences in AIc between sexes. To compare the distribution of individuals with negative or positive AIc values between sexes, we used the Fisher exact test in R 3.6.2. Additionally, since the dispersing sex includes both residents (with common genotypes) and immigrants (with rare genotypes), we expected a higher variance in AIc values for the more dispersing sex (Favre et al., 1997). An F-test for equality of variances was conducted to compare AIc variances between sexes in R 4.2.3 (R Core Team, 2024).

We also calculated the coefficient of relatedness between males and females of each species across the sampled localities using PLINK v2.0 (Purcell, 2007). We expect that the relatedness coefficient between same-sex individuals is influenced by dispersal bias dynamics. In this case, individuals of the non-dispersing sex are expected to have higher values of relatedness coefficient, reflecting a higher potential for inbreeding, while those of the dispersing sex are expected to exhibit lower relatedness values due to dispersal from their natal area in search of mates.

### Competitive phenotypes

To investigate whether competitive phenotypes predict patterns of philopatry and immigration within each sex for the two jacana species, we compared competitive traits between adult individuals with negative AIc values (likely recent immigrants) and those with positive AIc values (likely philopatric). These competitive phenotypes, including body mass, tarsus, and wing spur length measurements, have been associated with territoriality and reproductive success in other studies (Emlen & Wrege, 2004; Lipshutz 2017). Furthermore, these species differences are more pronounced between females than males, which do not differ in body mass (Lipshutz 2017). We predicted that individuals with larger competitive phenotypes would be more philopatric due to their enhanced territorial capabilities.

We used morphometric data on competitive traits from Lipshutz et al. (2017, 2019). Mass measurements were recorded in grams using a Pesola scale, while wing spur and tarsus measurements were taken from both the right and left sides to the nearest tenth of a millimeter using Avinet dial plastic calipers, and then averaged for each individual. To develop a combined metric of the competitive morphological phenotype of Jacanas, we conducted a principal component analysis (PCA) in R 3.2.2, summarizing body mass, tarsus length, and wing spur length. We conducted PCA analyses for each species and we retained a principal component (PC) score with an eigenvalue greater than 1, referred to as the morphological index PC1, which accounted for 82.1% and 73.4% of the variation in competitive morphology, for the Northern and Wattled jacanas, respectively (Table 1).

**Table 1.** Mean and standard error for morphological traits related to competitive phenotype and territoriality of both sexes of Northern Jacana and Wattled Jacana.

| Morphological trait | Northern Jacana |  |  | Wattled Jacana |  |  |
| --- | --- | --- | --- | --- | --- | --- |
|  | PC1 | Female | Male | PC1 | Female | Male |
| Eingvalue | 2.46 |  |  | 2.20 |  |  |
| Percent variation | 82.1 |  |  | 73.4 |  |  |
| Body mass (g) | 0.36 | 168 ± 2.7 | 102.6 ± 1.6 | 0.39 | 145.1 ± 3.8 | 100.6 ± 1.7 |
| Average spur (mm) | 0.31 | 13.8 ± 0.4 | 9.3 ± 0.2 | 0.30 | 10.3 ± 0.3 | 7.9 ± 1.6 |
| Tarsus length (mm) | 0.31 | 61.1 ± 0.4 | 55.4 ± 0.3 | 0.30 | 59.1 ± 0.5 | 54.3 ± 0.4 |
| Sample size |  | 35 | 62 |  | 26 | 46 |

## Results and Discussion

### Dispersal is male-biased in Northern Jacanas

In Northern Jacanas, we found several lines of evidence supporting male-biased dispersal and female philopatry. Population-level analyses revealed higher genetic structure in females compared to males across geographic distances (mean pairwise *F*_ST_: females 0.015, males 0.006; ANCOVA, p = 0.001; Figure 1b). This was further supported by individual-based AIc analysis, where females showed significantly higher AIc values (29.39) than males (-17.76; permutation test, p = 0.022; Figure 2a). Positive AIc values, as seen in females, indicate individuals with genotypes more commonly shared with others in the same locality, suggesting longer-term philopatry. Conversely, negative AIc values, as observed in males, indicate fewer common genotypes in their sampled locality, suggesting they may be recent immigrants or descendants of recent immigrants (Goudet et al., 2002). Additionally, the distribution of likely immigrant individuals differed significantly between sexes (F test, p = 0.05; Fisher’s exact test, p = 0.002). To our knowledge, this is the first genetic evidence of male-biased dispersal and female philopatry in species with a polyandrous mating system, in which females compete for mates and males provide parental care.

**Figure 2.**
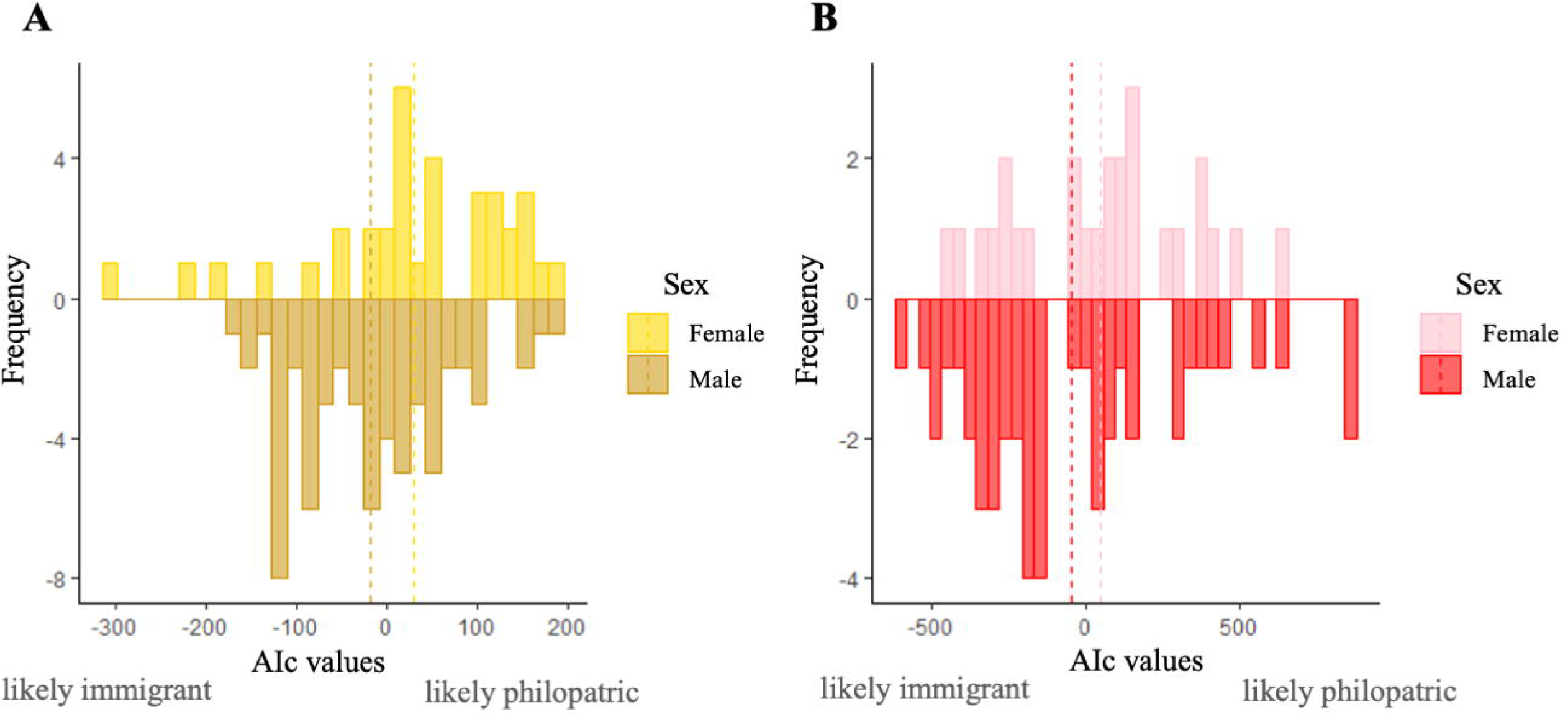
Assignment indices between sexes and *Jacana* species. Frequency distributions of the corrected assignment index (AIc) for females (bars above the axis) and males (bars below the axis) in (A) Northern Jacana and (B) Wattled Jacana. Positive AIc values suggest individuals are likely philopatric, meaning their genotype is common to others sampled from the same locality. Negative AIc values indicate likely recent immigrants or their descendants, with genotypes more closely related to individuals from different localities. Dashed lines show the mean AIc values.

In contrast, we found no statistical evidence of sex-biased dispersal in Wattled Jacanas. There were no significant differences in genetic structure across geographic distances (mean pairwise *F*_ST_: females 0.013, males 0.012; ANCOVA, p = 0.36; Figure 1c) nor AIc values (females 47.94, males -46.31; permutation test, p = 0.25; Figure 2b), nor the distribution of likely immigrants (F test, p = 0.22; Fisher’s exact test, p = 0.22). While these differences were not statistically significant, lower *F*_ST_ and negative AIc values for males in Wattled Jacanas suggest a potential trend toward male-biased dispersal.

Differences in the magnitude of sex-biased dispersal between Jacana species likely results from varying intensities of sexual selection. In Northern Jacanas, stronger sexual selection via female competition over mates and territories may drive the more pronounced female philopatry observed in our genetic results, compared to Wattled Jacanas. The sex ratio is a useful indicator of sexual selection pressure (Wade, 1979; Wade et al., 2003). Jacanas have a male-biased adult sex ratio, with many more males than females. Northern Jacana females have an average of ∼2.5 mates per territory, compared to ∼1.6 in Wattled Jacanas (Jenni & Collier, 1972; Emlen & Wrege, 2004). This higher number of mates in Northern Jacana suggests a greater degree of reproductive skew—where few individuals contribute disproportionately to the next generation (Shuster & Wade, 2003). Since detectable sex-biased dispersal rates with population genetics methods require relatively strong underlying pressures (Goudet et al., 2002), the stronger sexual selection in Northern Jacanas likely accounts for their stronger patterns of sex-biased dispersal.

A number of hypotheses have been proposed to link patterns of sex-biased dispersal with polygynous and socially monogamous mating systems, but previous examinations in polyandrous species have received mixed evidence for a clear pattern of sex-biased dispersal (Kwon et al., 2022). The local mate competition hypothesis has been predominantly supported in mammals with polygynous mating systems, in which male-biased dispersal is commonly found where males compete intensely for mates (Dobson, 1982). In birds with socially monogamous mating systems, the local resource competition hypothesis has been predominantly supported, for which female-biased dispersal is linked with male territorial defense (Greenwood, 1980). We find that female Northern Jacanas are more similar to socially monogamous males than to polygynous males, in that females are philopatric and males disperse. Thus, patterns of sex-biased dispersal in this polyandrous system are not simply the reverse of polygyny. Recent meta-analyses on sex-biased dispersal suggest that dispersal patterns are only weakly related to mating systems (Trochet et al., 2016; Végvári et al., 2018; Kwon et al., 2022), indicating that factors co-evolving with these systems, rather than the mating systems alone, influence genetic dispersal patterns.

In polyandrous systems, male-biased dispersal has been linked to the avoidance of kin competition, as males that disperse can reduce conflicts with relatives (Brom et al., 2016). Our relatedness data support this hypothesis, showing lower relatedness among males across sites compared to females in both Jacana species (Supplemental Material Figure S3). However, intense territorial competition among males may also influence this pattern, which could drive greater dispersal distances. While both sexes compete for territories, males in *Jacana* species are more aggressive as first responders to an intruder, and males compete with multiple co-mates for access to a single female partner (Emlen & Wrege, 2004; Lipshutz et al., 2017), potentially exacerbating dispersal as a result of disputes over territory. These dynamics may facilitate stronger dispersal patterns and, consequently, greater gene flow in males. Thus, male-biased dispersal in these systems could result from a combination of kin competition avoidance and territorial competition, highlighting the complex interplay of social and environmental factors in shaping dispersal patterns.

### Sexually selected phenotypes predict philopatry in female Northern Jacanas

We found a significant association between sexually selected, competitive phenotypes and philopatry in female Northern Jacanas, but not in Wattled Jacanas or males of either species (Figure 3). Specifically, philopatric Northern Jacana females exhibited significantly higher PC1 morphological index values compared to immigrant females (Figure 3a; p = 0.043), with these values primarily driven by body mass, followed by wing spur and tarsus length (Table 1). In contrast, Wattled Jacana females showed no significant differences in morphological values between philopatric and immigrant individuals (Figure 3b; p = 0.26), and males of both species also displayed no significant differences (p > 0.5).

**Figure 3.**
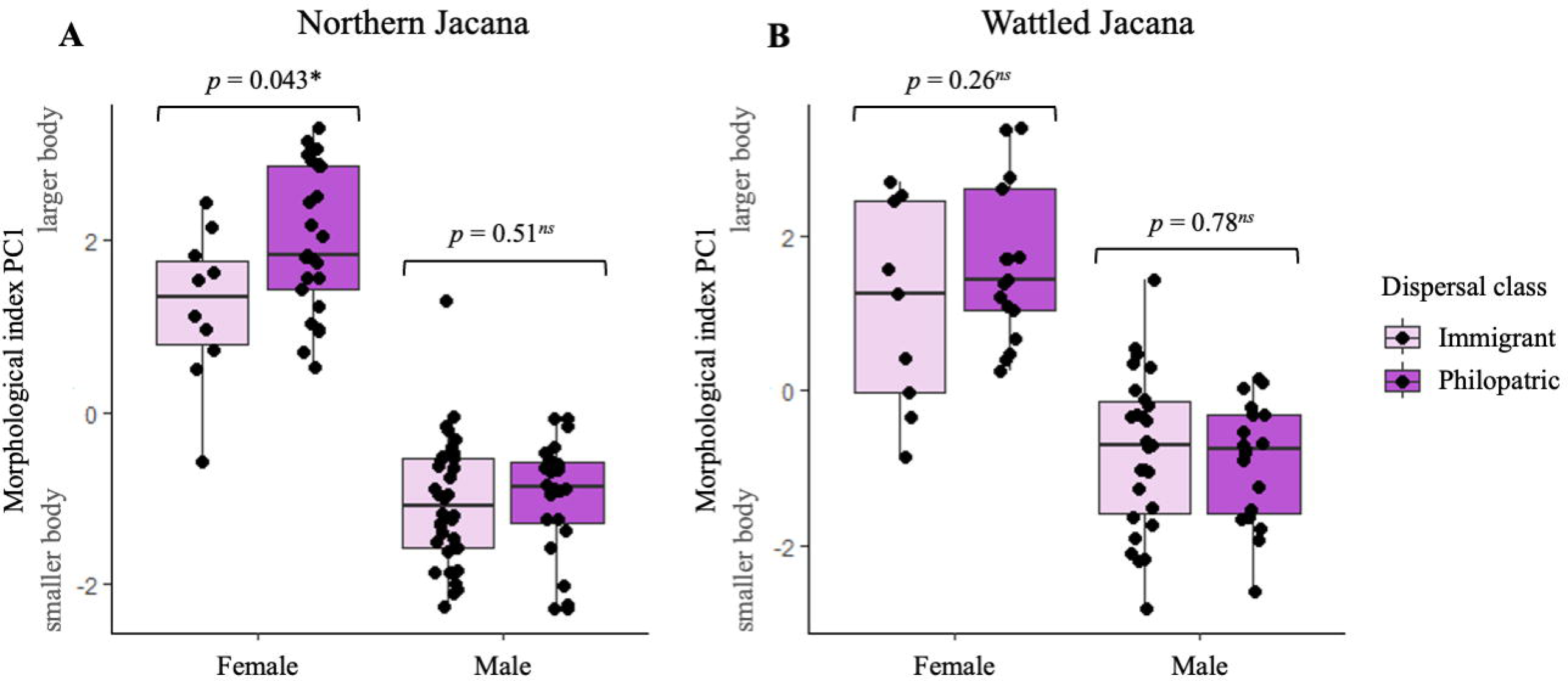
Comparison of competitive phenotype and dispersal classes. Non-parametric t-tests comparing PC1 morphological index values within each sex across dispersal categories (immigrant and philopatric) in (A) Northern Jacana and (B) Wattled Jacana. PC1 morphological index values represent combined body mass, wing spur size, and tarsus length metrics. Dispersal classes are derived from the assignment indices calculated for each individual. Symbols: * indicates *p* < 0.05, *ns* indicates non-significant p-values.

Sexually selected competitive phenotypes are critical predictors of territory and mate acquisition, which drive reproductive success across diverse mating systems (Arcese, 1989; Cassini, 2020). Sexual asymmetry in morphological traits, such as body and weapon size, may significantly influence sex-biased dispersal patterns across species (Trochet et al., 2016). The more competitive sex—typically males in polygynous systems—gains an advantage in maintaining a territory and competing for mates when individuals are larger (Candolin & Voigt, 2001; Cassini, 2020). Consequently, larger competitive traits can drive philopatry and limit gene flow, whereas smaller, less competitive individuals may disperse. In Jacanas, sexual dimorphism is female-biased, and more extreme in Northern Jacanas than in Wattled Jacanas (Lipshutz 2017). Our combined genetic and morphological results suggest that dispersal in Northern Jacana females is influenced by sexually selected phenotypes; in which individuals with larger body and weapon size are more philopatric and immigrant females are generally smaller (Figure 3a).

Ecological surveys of Northern Jacanas support this pattern, showing that larger females are more effective at maintaining territories (Jenni & Collier, 1972). This competitive advantage of body size in securing territories is also observed in other polyandrous species, such as the Spotted Sandpiper (Maxson & Oring, 1980), the Black-striped pipefish (Hubner et al., 2013) and bushcrickets (Schatral, 1993). This evidence supports the hypothesis that sexually selected traits, particularly competitive phenotypes, play a key role in shaping dispersal patterns across sexes (Trochet et al., 2016), influencing the spatial genetic structure of populations.

### Concluding remarks and perspectives

Due to their relative rarity in nature, and underrepresentation in the literature, polyandrous systems are seldom included in meta-analyses on sex-biased dispersal (e.g., Marbry et al., 2013; Trochet et al., 2016; Végvári et al. 2018). As a result, the general models for sex-biased dispersal tend to rely on data from more common mating systems, such as polygyny and social monogamy. Our results indicate that the selective pressures shaping male-biased dispersal in polygynous systems are not simply reversed in polyandrous systems. There are evolutionary pressures unique to polyandrous mating systems, such as the benefits females gain from multiple mates and the costs of egg production, which demand significant energy (Williams, 2005).

Rather than avoiding competition by dispersing (as is more typical in polygynous males), polyandrous females may prioritize staying to defend territories and attract mates. Consequently, it is plausible that, in polyandrous systems, female competition focuses more on resource defense than on active dispersal to find mates.

Jacanas are a classic example of social polyandry (Emlen & Oring, 1977), and as demonstrated here in our tests of sex-biased dispersal, they serve as an excellent model for testing hypotheses in the evolution of mating systems. Future work should examine a broader range of socially polyandrous species, combining population genetics with a phylogenetic comparative framework to determine whether the observed patterns of male-biased dispersal are unique to the Jacana species studied here or indicative of a more widespread phenomenon. Ultimately, these insights will enhance our understanding of the role of mating systems in structuring sex-biased dispersal within and among species. More broadly, the current evolutionary framework would benefit from additional focus on female perspectives of sexual selection (Ah-King, 2022). The evolution of competitive pressures in females is important not only for sex-biased dispersal, but also for female-female aggression (Cain & Ketterson, 2013), secondary sexual traits (Nalazco et al., 2022), and female-biased size dimorphism (Zamudio, 1998; Kilanowski et al., 2017). Integrating female competition into theoretical and empirical models will refine our understanding and interpretations of evolutionary patterns (Hare & Simmons, 2018; Fromonteil et al., 2023), enabling a more comprehensive framework that accounts for diverse selective pressures across both sexes and different mating systems.

## Data availability statement

Genotype-by-sequencing data can be found on NCBI under BioProject PRJNA494934 and BioSamples SAMN10185940–SAMN10186205. The raw and filtered VCF files, morphological data, as well as the software codes, are available in the Dryad Digital Repository at DOI: xxxxx.

## Acknowledgements

We thank the Department of Biology at Duke University for providing facilities and administrative support. We are also grateful to our colleagues Gabriel Macedo, Tessa Paton, Valentina Gómez-Bahamón, and Kevin Bennett for their helpful feedback on the initial manuscript draft, and to Jordan Karubian for inspiring this study. Bioinformatics analyses for this research were conducted on Duke University’s high-performance Compute Cluster.

## Author contributions

Study conceptualization, L.W.L., S.E.L., Data collection, S.E.L., Computation and statistical analysis, L.W.L., Figure Visualization, L.W.L., Manuscript writing, review, and editing, L.W.L., S.E.L.

