## Supplementary material for "Sexually selected traits predict female-biased philopatry in a sex-role reversed shorebird": Online Supplemental Information

**Table of contents**


| | |
|---|---|
| Proportion of genetic admixture | Page 2 |
| Classification of hybrids and parental genotypes | Page 3 |
| Relatedness coefficient by locality between males and females of both species | Page 4 |

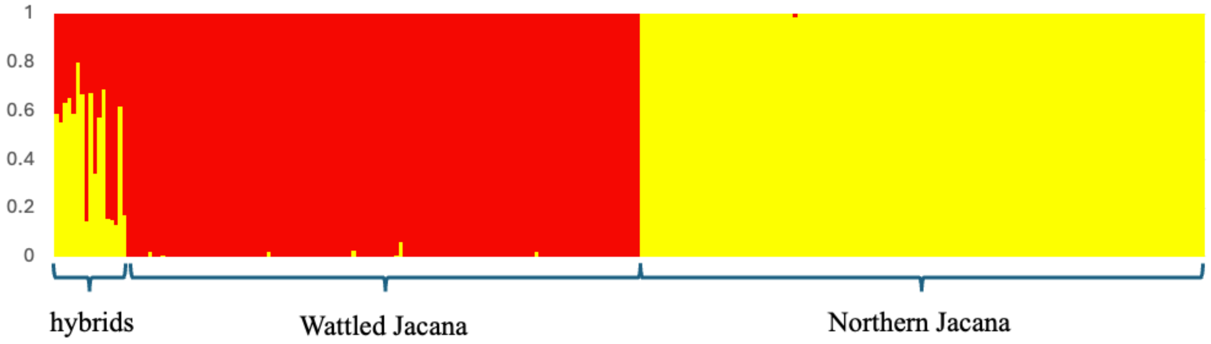


**Figure S1.** Proportion of genetic admixture in hybrids and parentals of Northern Jacana and Wattled Jacana. Individuals were classified as putative hybrids if their admixture proportion was greater than 0.1 and less than 0.9.

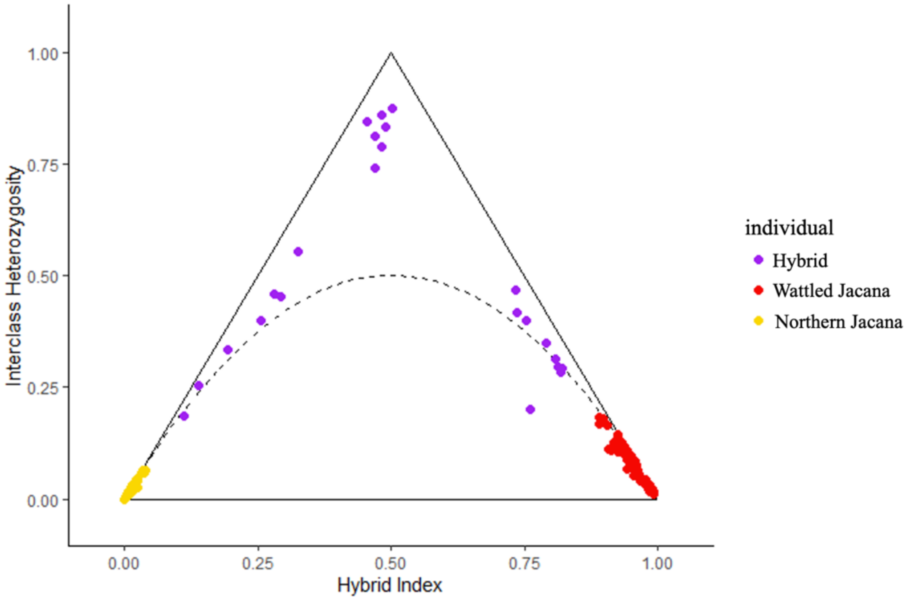


**Figure S2.** Classification of hybrids and parental genotypes, showing F1 hybrids and backcrosses (purple dots), along with Northern Jacana parentals (yellow dots on the left corner) and Wattled Jacana parentals (red dots on the right corner).

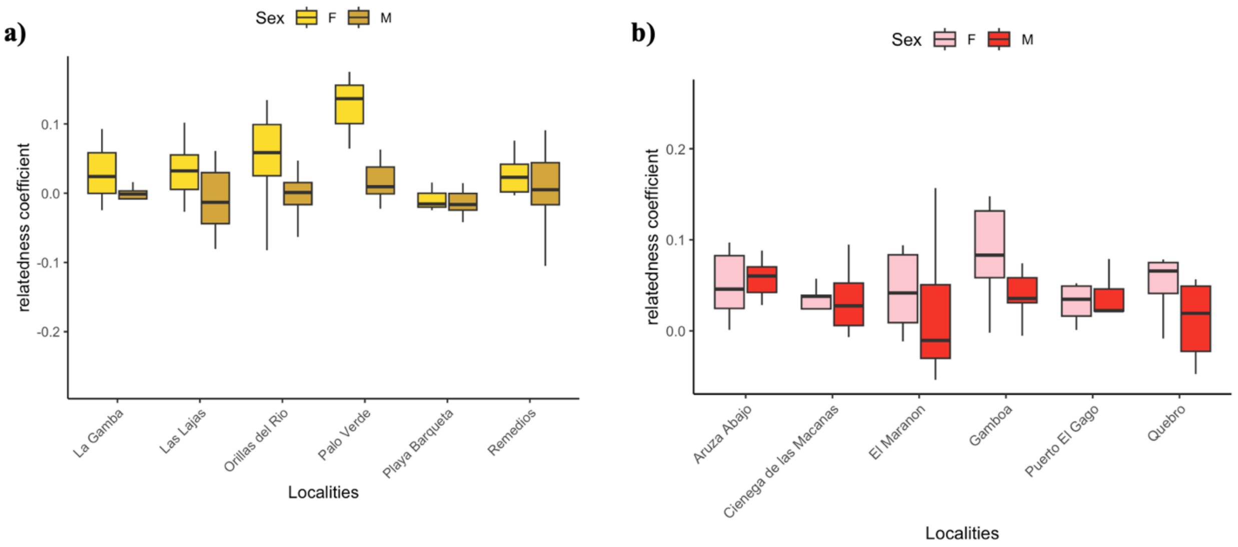


**Figure S3.** Relatedness coefficient by locality between males and females of (a) Northern Jacana and (b) Wattled Jacana.
